# Composition of a low erucic acid, low fiber field pennycress (*Thlaspi arvense* L) grain referred to as CoverCress™ developed through breeding and gene editing

**DOI:** 10.1101/2022.06.03.494728

**Authors:** Gary Hartnell, Shawna Lemke, Chris Aulbach

## Abstract

Field pennycress (*Thlaspi arvense* L.) can be domesticated and cultivated as an annual in a corn / pennycress / soybean rotation where pennycress is sown and harvested as a cash cover crop. To improve the nutritional profile, pennycress was modified in two ways to achieve the same alterations in characteristics: 1) through selection of mutants and 2) through gene editing. These alterations resulted in a low erucic acid, lower fiber phenotype and the resulting products from these combinations are referred to as CoverCress CCWG-1 and CoverCress CCWG-2, respectively. CCWG-1 and CCWG-2 were planted as cover crops in five U.S. locations in the fall and the grain was harvested in the subsequent June. The grain was treated with 83 mM copper sulfate solution as a potential palatability agent for the naturally high glucosinolate levels, and was subsequently analyzed for nutrient (proximates, minerals, fatty acids, amino acids, and vitamins) and anti-nutrient (sinapine, glucosinolates, mold) content. The low erucic acid, lower fiber phenotype was consistently achieved across five lots. Generally, the nutrient content for both CCWG-1 and CCWG-2 were similar to canola grain. Canola grain contains the anti-nutrient sinapine but is significantly reduced to below the level of detection in CoverCress grain. As expected, CoverCress grain contains about 10 times more glucosinolates than canola. Based on the composition of CoverCress grain, it may provide a source of energy and amino acids to animals with restricted inclusion based on the glucosinolate content.

## Introduction

Field pennycress (*Thlaspi arvense L.*) is an annual weed that grows over the winter in Eurasia and North America. It is a member of the Brassicacae family, commonly known as the mustard family. Field pennycress contains high levels of oil (~25-30%) that makes it a desirable ultra-low carbon fuel feedstock. The potential commercial value of field pennycress production has been researched extensively in a variety of leading laboratories including the National Center for Agricultural Innovation, USDA ARS, Peoria, IL, Western Illinois University, Illinois State University and the University of Minnesota. The cumulative literature and experience establish that field pennycress can be domesticated and cultivated as a winter annual in a corn / pennycress / soybean rotation where pennycress is sown and harvested as a cash winter cover crop as (Phippen and Phippen, 2012; Sedbrook *et al*., 2014).

In addition to this primary value for fuel, the seed could provide an energy source for animal feeds such as chicken feed, however, the oil typically contains >35% erucic acid. Erucic acid is a 22-carbon monounsaturated acid that is absorbed, distributed and metabolized like other fatty acids involving primarily metabolism via mitochondrial beta-oxidation and, to a lesser extent, peroxisomal beta-oxidation. Like other longer-chain fatty acids, the rate of mitochondrial beta oxidation is comparatively lower for erucic acid; however, elevated erucic acid levels induce liver peroxisomal oxidation pathways as a mechanism of compensation. Interest in the safety of erucic acid occurred when results of studies in rats associated the dietary intake of high doses of erucic acid with myocardial lipidosis and heart lesions (Bremer and Norum, 1982). Oilseed rape conventionally contains similarly high levels of erucic acid. Low erucic acid varieties were identified and marketed as canola, which have been shown to be safe to broiler chickens when incorporated in diets up to 25% inclusion. Reduction in erucic acid is achieved through disruption of Fatty Acid Elongation 1 (FAE1), resulting in higher levels of oleic (18:1). It is through this same mechanism that erucic acid levels has been lowered in pennycress (McGinn et al., 2019).

Field pennycress is also high in fiber, which can impact digestibility as a feed ingredient. The production of seed coat fiber was first characterized in the model plant *Arabidopsis. Arabidopsis* seed coats derive their brown color from the accumulation of proanthocyanidins (PAs), a class of flavonoid chemicals (polymerized flavan-3-ols, or condensed tannins) that protect against a variety of biotic and abiotic stresses and help maintain seed dormancy and viability (Debeaujon *et al*., 2003). PAs start out as colorless epicatechin compounds until they are transported to the vacuole where they are polymerized and oxidized as the seed desiccates. In *Arabidopsis*, PAs are only produced in a narrowly defined cell layer in the endothelium of the seed, and the genes TTG1, TT8/bHLH042, and TT2/MYB123 and have been demonstrated as being the three main regulators of PA biosynthesis in seed coat (Baudry *et al*., 2004; Lepiniec *et al*., 2006). Gonzalez *et al*. (2009) described how the TTG1 works in a complex with a particular combination of MYB class and bHLH class transcription factors to regulate epidermal development of the seed coat. Loss-of-function mutants in these genes exhibit the transparent “testa” phenotype as a result of low levels of oxidized PAs in the seed coat. Similarly, the transparent testa phenotype was observed with loss-of-function mutations in orthologs of these genes in pennycress (Chopra *et al*, 2018), resulting in reduced fiber content. This phenomenon has been observed in other brassicas such as canola and is characterized by yellow seeds that have more oil because of the resulting thinner seed coat and larger embryo (Abraham & Bhatia, 1986). Meal from these brassicas have also been shown to be useful in animal feed because of the relatively lower fiber and higher metabolizable energy (Slominski *et al*., 1994, Slominski, 2015;).

Reduction of erucic acid and fiber levels in pennycress has been achieved through conventional breeding and gene editing for loss of function in the associated gene pathways. A low-erucic acid, lower-fiber pennycress is being developed under the trade name of CoverCress™.

CoverCress^1^ represents a clear opportunity for sustainable optimization of certain agricultural systems. It serves as an important winter cover, working within the no/low-till cropping systems to prevent soil erosion from fallow fields and improve soil nitrogen management. As part of the safety assessment of this new grain for animal feed, a comprehensive compositional study was conducted on both the mutant breeding line and gene edited lines.

## Materials and Methods

### CoverCress Line Development

Two studies were conducted to assess the composition of CoverCress grain. The first study utilized grain from a low-erucic acid, lower-fiber variety of CoverCress developed through breeding by identification of mutants. One parent line, referred to as MN106-V300, was developed via ethyl methanesulfonate (EMS) mutation breeding. This line contains a mutation to the FAE1 gene (specifically, a cytosine to thymine change at base position 1018bp), resulting in a premature stop codon and complete loss of function of the gene, causing a reduction in erucic acid level (Chopra *et al*., 2020). The second parent line, referred to as Y1126, was isolated from a cultivated field in Grantfork IL and was identified as a lower-fiber phenotype with a yellow seed coat. This line contains a naturally-occurring deletion of 21bp in the TRANSPARENT TESTA GLABRA 1 (TTG1) gene. This mutation results in a deletion of 7 amino acids in the conserved area of TTG1 protein, leading to a complete loss of function. The MN106-V300 and Y1126 lines were crossed in the fall, and F2 plants carrying homozygous mutant alleles for these genes were further propagated in bulk, first in local greenhouses in the spring and then in the field in an 0.3-acre increase planted at Sigel, IL in the subsequent fall. Seeds from this increase were harvested in bulk in the spring, and this seed formed the foundation source. It should be noted that this source is segregating for all background genetics that differ between Y1126 and MN106-V300 and fixed only for FAE1 and TTG1. This line is referred to as CoverCress Whole Grain (CCWG-1).

The second study used a line that was generated through application of gene editing techniques to resulting in lines of low-erucic acid, lower-fiber pennycress and is referred to as CCWG-2. To generate this line, mutations were introduced into a pennycress cultivar using a CRISPR/SpCas9 DNA construct designed to target genomic edits to the FAE1 and TT8 genes. This transgene construct was delivered to a pennycress cultivar using a disarmed Agrobacterium tumefaciens strain (GV3101) and a standard floral dip transformation method. Presence of the edits in T1 plants was confirmed through multiple methods including confirmatory PCR screening of a fragment of the T-DNA. Seed from the progeny T2 generation was screened for segregants that did not have the transgene.

Resulting progeny in the T3 generation were screened again for negative presence of DsRED and Cas as well as homozygous edits to TT8 and FAE1. The resulting edited plants contain small indels in both the FAE1 and TT8 genes that lead to loss of function due to premature stop codons. The edited plants produced yellow seed (a marker for low fiber) and low accumulation of erucic acid in seeds and were taken forward for subsequent characterization and seed bulk up.

### Grain production

Two studies were conducted to measure the nutrient and antinutrient content of CoverCress grain. In Study A, CCWG-1 was grown in five locations (Venedy, IL; Sigel, IL; Havana, IL; and two locations in Mt. Pulaski, IL). In Study B, CCWG-2 was grown in five locations (Havana, IL; Arenzville, IL; Mt. Pulaski, IL: and two locations in Martinsville, IL).

### Treatment of grain with copper sulfate

Native pennycress and CoverCress grain contain high levels of glucosinolates in the form of aliphatic sinigrin (approximately 35-110 μ moles/g meal). Glucosinolates are considered an antinutrient and can prevent the absorption of iodine and may affect thyroid function in target species (Schöne et al., 1993). Glucosinolates can also impart a bitter flavor which may affect feed consumption (Bischoff, 2021). Copper sulfate treatment has been suggested to overcome the deleterious effects of glucosinolates for other high glucosinolate grains (Schöne *et al*., 1993; Singhal & Sinha 2000; Payvastagan, 2013). It was theorized that copper may reduce or mask the bitter taste of glucosinolates, improving palatability. Therefore, both the CCWG-1 and CCWG-2grains were pre-treated with copper sulfate prior to characterization of the grain composition and use in subsequent broiler feeding studies.

Three hundred grams of CCWG-1 grain from each of the five locations was blended with 45 ml of 83.3 mM copper sulfate pentahydrate solution and referred to as CCWG-1-CuSO_4_. Four hundred and sixty grams of CCWG-2 grain from each of the five locations were blended with 60 ml of 83.3 mM copper sulfate pentahydrate solution and referred to as CCWG-2-CuSO_4_.. Samples were stored at 2 to 8°C.

### Methods of Analysis

The five lots of CCWG-1-CuSO_4_ and CCWG-2-CuSO_4_ were analyzed in duplicate by EPL Bio Analytical Services (Niantic, IL) following Good Laboratory Practices (GLP) and by Dairyland Laboratories (Arcadia, WI) a commercial laboratory that uses methods on the latest research findings and technical methodologies available to the feed industry. Dairyland Laboratories participates (DLL) in the National Forage Testing Association, North American Proficiency Testing Program (NAPT), and American Association of Feed Control Officials (AAFCO) Proficiency Testing Program. These programs help laboratories generate accurate and precise analyses.

The list of analytes and the respective Standard Operating Procedure (SOP) used by each lab are shown in Table 1. The details of the specific analytical methodology used can be obtained from the respective laboratories.

**Table 1.** List of analytes measured by EPL Bio Analytical Services (EPL) and Dairyland Laboratories and Standard Operating Procedures (SOP).

| Analyte | EPL | Dairyland Laboratories |
| --- | --- | --- |
|  | SOP | SOP |
| Moisture/Dry Matter | GSOP-NC-4 | DLARC.SOP.FEED.WCFEED.Lab Dry Matter 105<br>3hr.001 |
| Crude Protein | GSOP-NC-20 | DLARC.SOP.FEED.WCFEED.Leco528.008 |
| Ash | GSOP-NC-2 | DLARC.SOP.FEED.WCFEED.Ash.003 |
| Crude Fiber | GSOP-NC-5 | DLARC.SOP.FEED.WCFEED.Crude Fiber.006 |
| Acid Detergent Fiber (ADF) | GSOP-NC-3 | DLARC.SOP.FEED.WCFEED.Sequential<br>NDF/ADF/Lignin Analysis.000 |
| Neutral Detergent Fiber (NDF) | GSOP-NC-9 | DLARC.SOP.FEED.WCFEED.Sequential<br>NDF/ADF/Lignin Analysis.000 |
| Total Dietary Fiber (TDF) | GSOP-NC-359 | Not measured |
| Crude Fat (Acid hydrolysis) | GSOP-NC-141 | DLARC.SOP.FEED.WCFEED.AH Fat.002 |
| Minerals (Ca, P, Mg, K, Na, S,<br>Cu, Mn, Fe, Zn) | GSOP-NC-60 | DLARC.SOP.FEED.WCFEED.ICP Analysis.006 |
| Chloride | GSOP-NC-328 | DLARC.SOP.FEED.WCFEED.Salt.006 |
| Carbohydrates | GSOP-NC-494 | Not Measured |
| Cysteine and Methionine | GSOP-NC-279 | DLARC.SOP.FEED.WCFEED.Cysteine and<br>Methionine.000 |
| Tryptophan | GSOP-NC-22 | DLARC.SOP.FEED.WCFEED.Tryptophan Analysis.001 |
| 15 Amino Acids | GSOP-NC-58 | DLARC.SOP.FEED.WCFEED. Amino Acid<br>Analysis.000 |
| Fatty Acids | GSOP-NC-319 | DLARC.SOP.FEED.WCFEED.Fatty Acid Analysis.004 |
| Vitamin B1 (Thiamine) | GSOP-NC-262 | Not Measured |
| Vitamin B2 (Riboflavin) | GSOP-NC-46 | Not Measured |
| Vitamin B6 (Pyridoxine) | GSOP-NC-220 | Not Measured |
| Folic Acid | GSOP-NC-26 | Not Measured |
| Niacin | GSOP-NC-28 | Not Measured |
| Tocopherols (Alpha, beta,<br>gamma, delta) | GSOP-NC-341 | Not Measured |
| Fumonisin (B1, B2, B3) | GSOP-NC-336 | Not Measured |
| Aflatoxin (B1, B2, G1, G2), T-2 Toxin, Ochratoxin-A | GSOP-NC-337 | Not Measured |
| Deoxynivalenol, 3/15-acetyl-deoxynivalenol, Zearalenone | GSOP-NC-338 | Not Measured |
| Ergosine, Ergotamine, Ergocornine, Ergocryptine, Ergocristine | GSOP-NC-456 | Not Measured |
| Sinapine | GSOP-NC-208 | Not Measured |
| Sinigrin | GSOP-NC-650 | Not Measured |

### Data Handling

The data for each study consisted of five samples each from one of five locations grown in Illinois with duplicate values for each sample. The duplicates were averaged to give one value for each of the five samples.

For the data generated by EPL, Statistical Consultants Plus, LLC (Fenton, MO) calculated measures of data dispersion (variability/standard deviation and data ranges) and measures of central tendency (mean and median) for each analyte using the analytical data from the five test substances each representing five different field locations using SAS, version 9.4. For the data generated by Dairyland Laboratories, Hartnell International Consulting LLC (St. Peters, MO) calculated measures of data dispersion (variability/standard deviation and data ranges) and measures of central tendency (mean and median) for each analyte. Measures were calculated using Microsoft Excel.

## Results and Discussion

### Study A – CCWG-1-CuSO_4_

#### Proximates

Table 2 contains the results of proximate analyses. In general, data were in agreement between laboratories except for acid detergent fiber (ADF), neutral detergent fiber (NDF) and crude fat. The two labs used different fat extraction methods. Methodology was also different for the analysis of ADF and NDF, however, in both cases the ADF and NDF analyses were conducted on the residue after fat extraction. It is postulated that Dairyland Laboratories’s method extracted more fat from the product than EPL’s method and that EPL’s assessment of ADF and NDF may be higher because of residual fat, which inflated the ADF and NDF values. In addition, there was greater variation noted with EPL’s fat values than with Dairyland Labs. The Fiber Best Practices Working Group under AAFCO’s Laboratory and Services Committee provides a detailed discussion on the critical factors in determining fiber in feed and forages (AAFCO, 2017). As expected, mean moisture levels were 15.6 −17.1% because of the addition of the copper sulfate solution to the CoverCress grain. Mean and median values are similar for all parameters, indicating that the values from the five lots are evenly distributed, i.e. the data are symmetrical in distribution.

**Table 2.** CCWG-1-CuSO_4_ Proximates (100% dry matter basis)

| Analytical |  | Standard |  |  |  | Range |  |
| --- | --- | --- | --- | --- | --- | --- | --- |
| Lab<sup>1</sup> | Variable | Units | Mean | Deviation | Median | Minimum | Maximum |
| EPL | Moisture | % | 17.1 | 0.23 | 17.0 | 16.9 | 17.4 |
| DLL | Moisture | % | 15.6 | 0.34 | 15.5 | 15.1 | 16.1 |
| EPL | Dry Matter | % | 82.9 | 0.23 | 83.1 | 82.7 | 83.1 |
| DLL | Dry Matter | % | 84.4 | 0.34 | 84.5 | 83.9 | 84.9 |
| EPL | Crude Fiber | % | 23.1 | 1.28 | 22.8 | 21.5 | 25.0 |
| DLL | Crude Fiber | % | 23.0 | 1.72 | 22.7 | 20.9 | 24.9 |
| EPL | Acid Detergent Fiber | % | 30.1 | 1.54 | 29.9 | 28.0 | 32.1 |
| DLL | Acid Detergent Fiber | % | 13.9 | 1.58 | 13.5 | 12.1 | 15.8 |
| EPL | Neutral Detergent Fiber | % | 29.5 | 1.55 | 29.2 | 27.7 | 32.0 |
| DLL | Neutral Detergent Fiber | % | 20.3 | 2.76 | 19.3 | 17.3 | 24.5 |
| EPL | Total Dietary Fiber | % | 30.0 | 0.97 | 29.8 | 28.9 | 31.5 |
| EPL | Crude Protein | % | 25.1 | 1.53 | 25.5 | 22.6 | 26.7 |
| DLL | Crude Protein | % | 25.5 | 1.63 | 25.6 | 22.7 | 26.7 |
| EPL | Crude Fat | % | 32.7 | 1.26 | 33.1 | 30.6 | 33.7 |
| DLL | Crude Fat | % | 36.3 | 0.52 | 36.5 | 35.5 | 36.7 |
| EPL | Carbohydrates | % | 36.9 | 1.61 | 36.9 | 34.6 | 38.5 |
| EPL | Ash | % | 5.3 | 0.39 | 5.5 | 4.9 | 5.7 |
| DLL | Ash | % | 5.8 | 0.40 | 5.9 | 5.3 | 6.2 |

<sup>1</sup> Analytical Lab: EPL – EPL Bio Analytical Services; DLL – Dairyland Laboratories

#### Amino Acid analysis

Table 3 contains the amino acid profile of CCWG-1-CuSO_4_ when expressed on a 100% dry matter basis and Table 4 contains the amino acid profile of CCWG-1-CuSO_4_ when expressed on a % of total protein basis. The amino acid values from the five lots showed a symmetrical distribution with the means and median values being similar. EPL consistently had lower variability (smaller standard deviations) than Dairyland Laboratories. Dairyland reported numerically higher values for most amino acids whether expressed as a percent of dry matter or as a percent of protein. Dairyland had a higher percentage of total amino acids when expressed as total amino acids as a percent of crude protein (97% versus 89.4%).

**Table 3.**
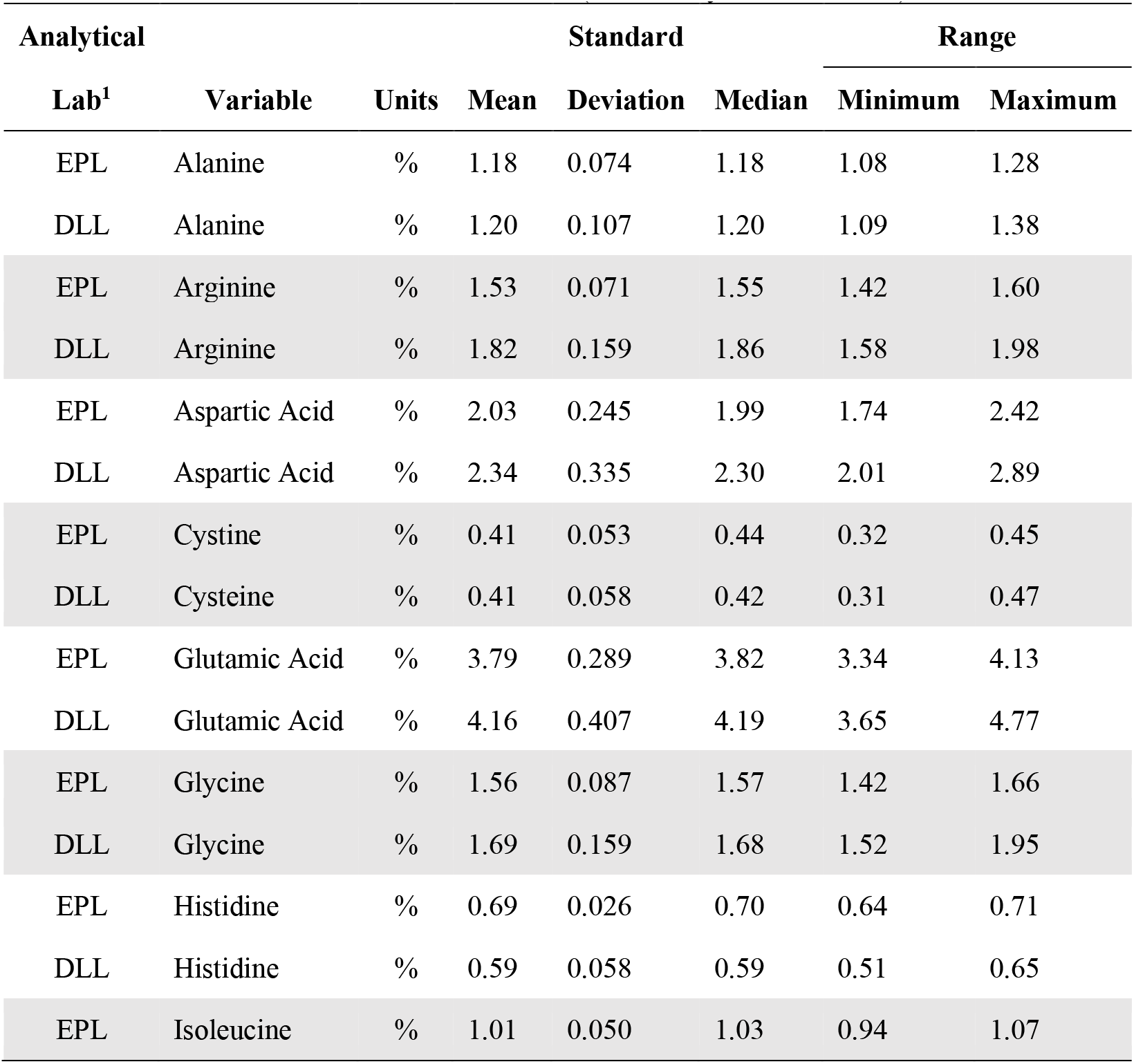
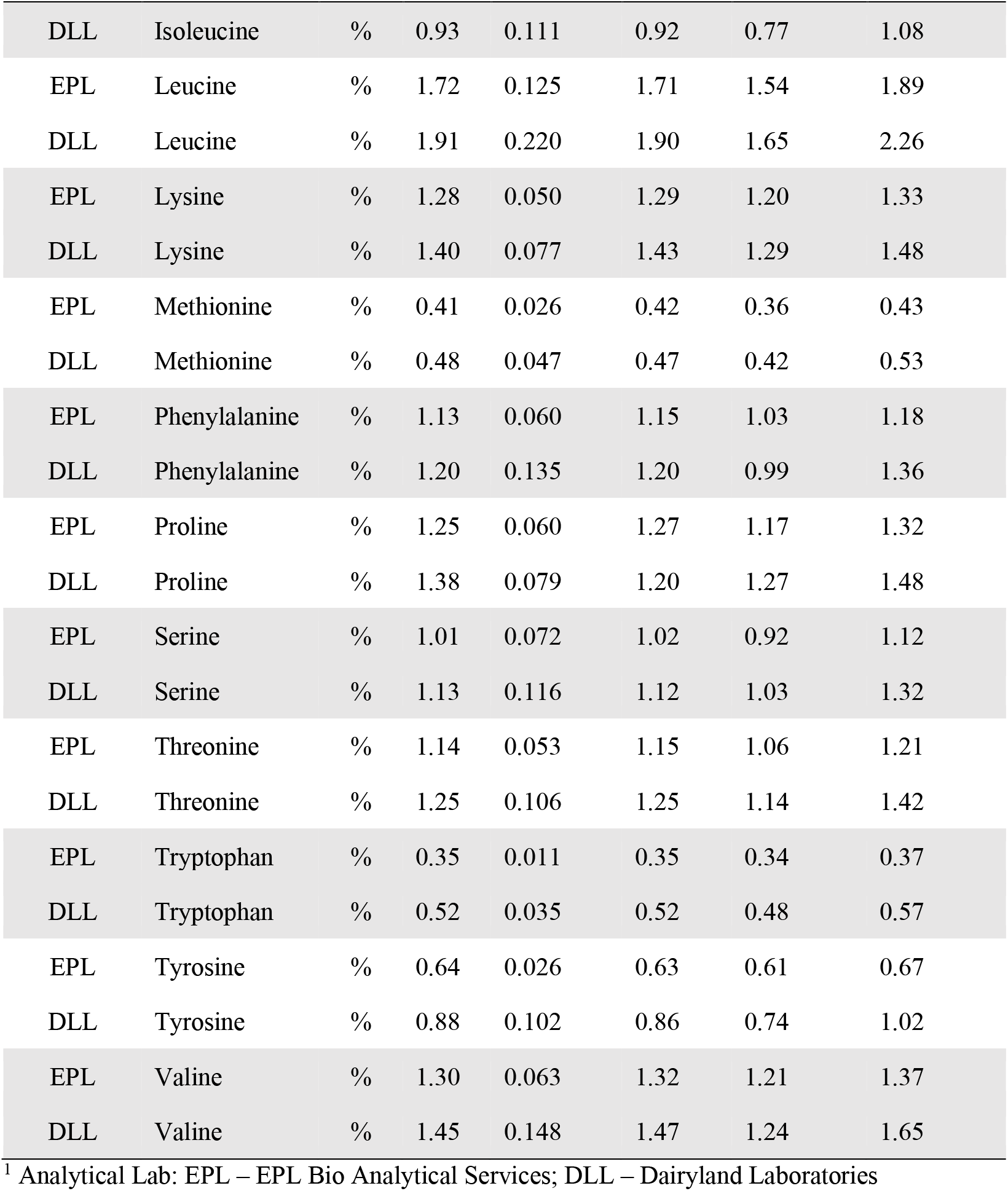
CCWG-1-CuSO_4_ Amino Acids (100% dry matter basis)

| Analytical |  | Standard |  |  |  | Range |  |
| --- | --- | --- | --- | --- | --- | --- | --- |
| Lab<sup>1</sup> | Variable | Units | Mean | Deviation | Median | Minimum | Maximum |
| EPL | Alanine | % | 1.18 | 0.074 | 1.18 | 1.08 | 1.28 |
| DLL | Alanine | % | 1.20 | 0.107 | 1.20 | 1.09 | 1.38 |
| EPL | Arginine | % | 1.53 | 0.071 | 1.55 | 1.42 | 1.60 |
| DLL | Arginine | % | 1.82 | 0.159 | 1.86 | 1.58 | 1.98 |
| EPL | Aspartic Acid | % | 2.03 | 0.245 | 1.99 | 1.74 | 2.42 |
| DLL | Aspartic Acid | % | 2.34 | 0.335 | 2.30 | 2.01 | 2.89 |
| EPL | Cystine | % | 0.41 | 0.053 | 0.44 | 0.32 | 0.45 |
| DLL | Cysteine | % | 0.41 | 0.058 | 0.42 | 0.31 | 0.47 |
| EPL | Glutamic Acid | % | 3.79 | 0.289 | 3.82 | 3.34 | 4.13 |
| DLL | Glutamic Acid | % | 4.16 | 0.407 | 4.19 | 3.65 | 4.77 |
| EPL | Glycine | % | 1.56 | 0.087 | 1.57 | 1.42 | 1.66 |
| DLL | Glycine | % | 1.69 | 0.159 | 1.68 | 1.52 | 1.95 |
| EPL | Histidine | % | 0.69 | 0.026 | 0.70 | 0.64 | 0.71 |
| DLL | Histidine | % | 0.59 | 0.058 | 0.59 | 0.51 | 0.65 |
| EPL | Isoleucine | % | 1.01 | 0.050 | 1.03 | 0.94 | 1.07 |
| DLL | Isoleucine | % | 0.93 | 0.111 | 0.92 | 0.77 | 1.08 |
| EPL | Leucine | % | 1.72 | 0.125 | 1.71 | 1.54 | 1.89 |
| DLL | Leucine | % | 1.91 | 0.220 | 1.90 | 1.65 | 2.26 |
| EPL | Lysine | % | 1.28 | 0.050 | 1.29 | 1.20 | 1.33 |
| DLL | Lysine | % | 1.40 | 0.077 | 1.43 | 1.29 | 1.48 |
| EPL | Methionine | % | 0.41 | 0.026 | 0.42 | 0.36 | 0.43 |
| DLL | Methionine | % | 0.48 | 0.047 | 0.47 | 0.42 | 0.53 |
| EPL | Phenylalanine | % | 1.13 | 0.060 | 1.15 | 1.03 | 1.18 |
| DLL | Phenylalanine | % | 1.20 | 0.135 | 1.20 | 0.99 | 1.36 |
| EPL | Proline | % | 1.25 | 0.060 | 1.27 | 1.17 | 1.32 |
| DLL | Proline | % | 1.38 | 0.079 | 1.20 | 1.27 | 1.48 |
| EPL | Serine | % | 1.01 | 0.072 | 1.02 | 0.92 | 1.12 |
| DLL | Serine | % | 1.13 | 0.116 | 1.12 | 1.03 | 1.32 |
| EPL | Threonine | % | 1.14 | 0.053 | 1.15 | 1.06 | 1.21 |
| DLL | Threonine | % | 1.25 | 0.106 | 1.25 | 1.14 | 1.42 |
| EPL | Tryptophan | % | 0.35 | 0.011 | 0.35 | 0.34 | 0.37 |
| DLL | Tryptophan | % | 0.52 | 0.035 | 0.52 | 0.48 | 0.57 |
| EPL | Tyrosine | % | 0.64 | 0.026 | 0.63 | 0.61 | 0.67 |
| DLL | Tyrosine | % | 0.88 | 0.102 | 0.86 | 0.74 | 1.02 |
| EPL | Valine | % | 1.30 | 0.063 | 1.32 | 1.21 | 1.37 |
| DLL | Valine | % | 1.45 | 0.148 | 1.47 | 1.24 | 1.65 |

<sup>1</sup> Analytical Lab: EPL – EPL Bio Analytical Services; DLL – Dairyland Laboratories

**Table 4.**
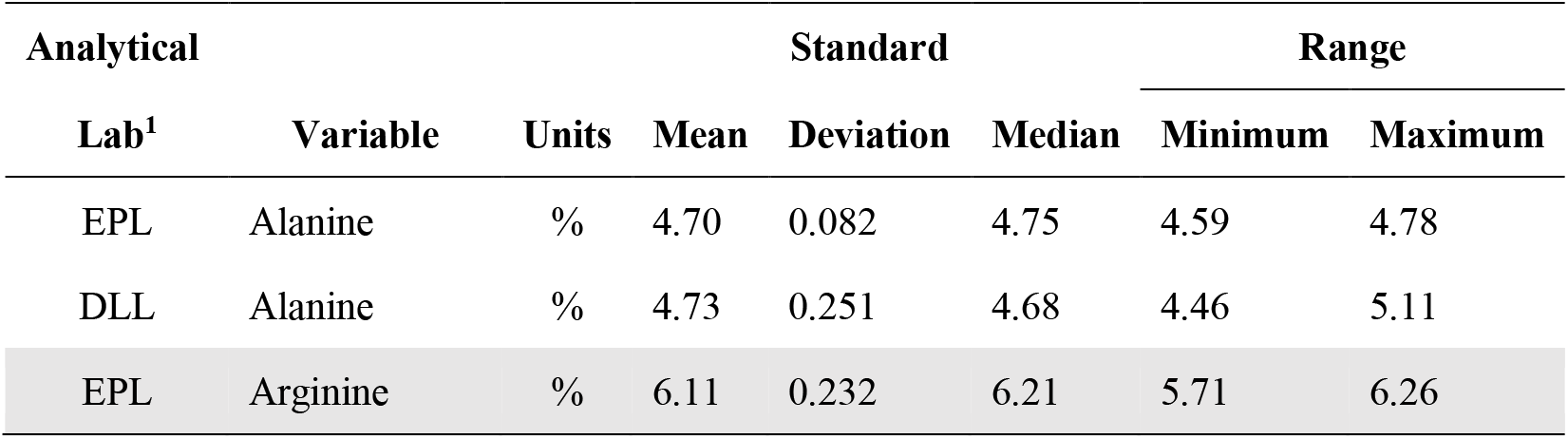
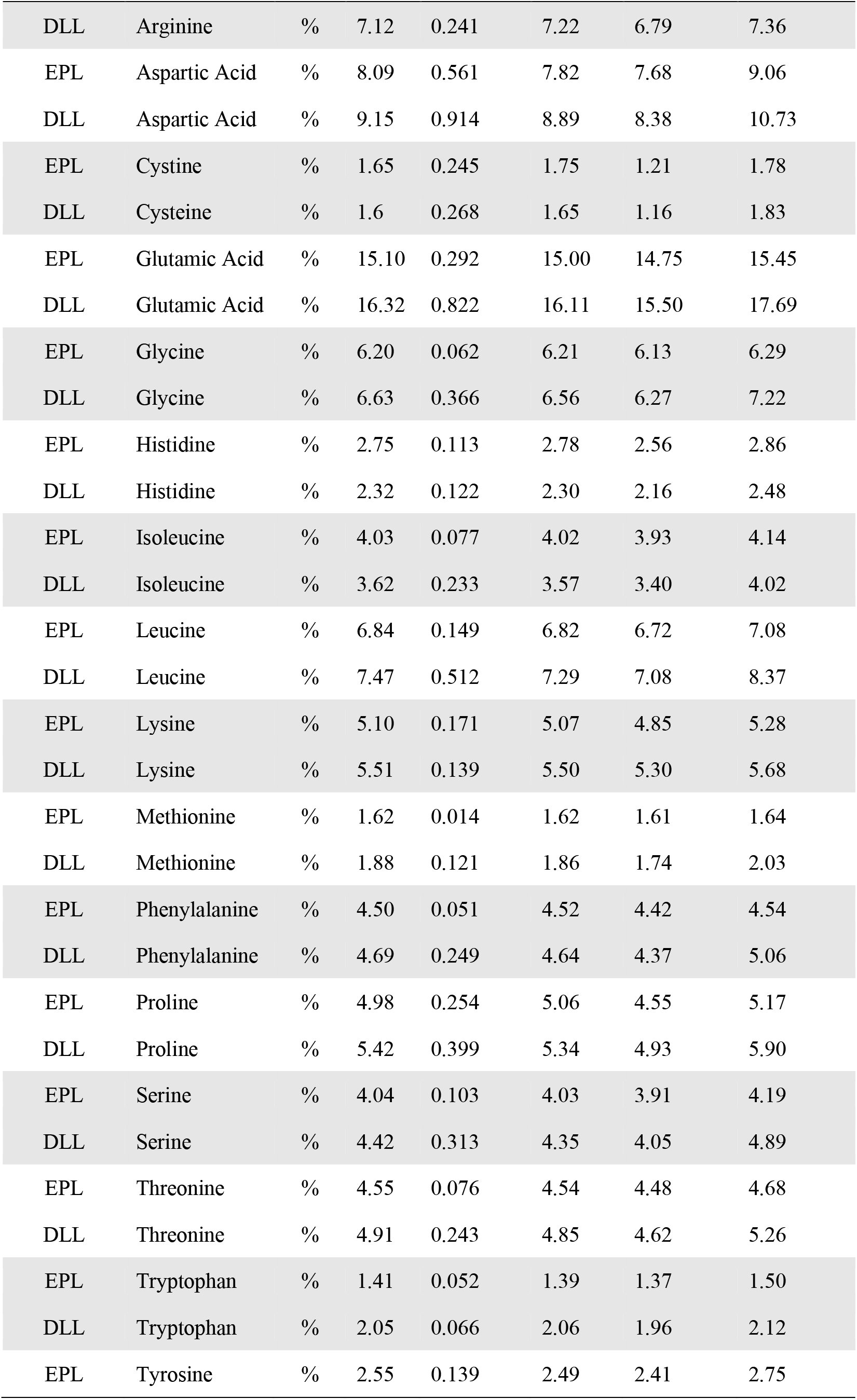
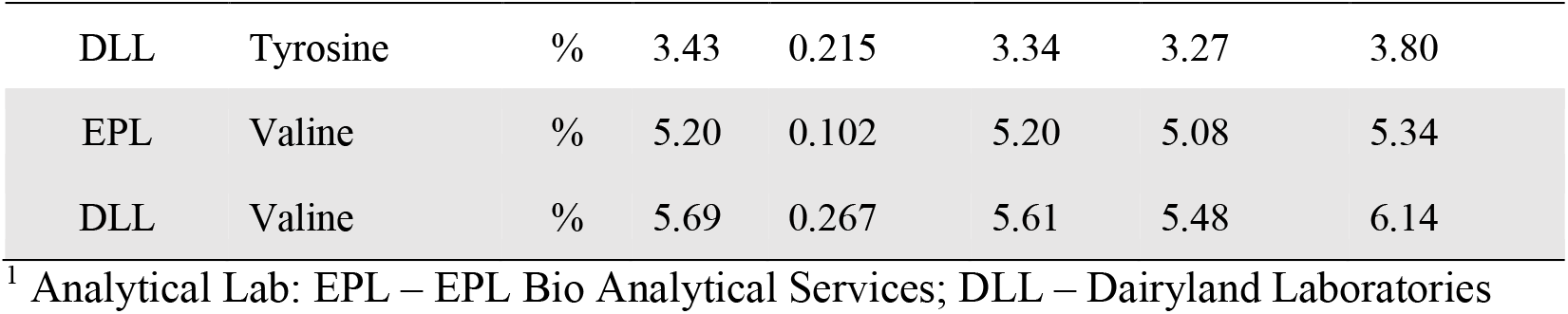
CCWG-1-CuSO_4_ Amino Acids (% of protein basis)

#### Fatty Acid Characterization

Tables 5 and 6 contain the fatty acid profile of CCWG-1-CuSO_4_ when expressed on a 100% dry matter basis and percent of total fatty acids, respectively. In general, there was good agreement between laboratories. The fatty acid values from the five lots showed a symmetrical distribution with the means and median values being similar. The major fatty acids are oleic (40% of the fatty acids) followed by linoleic (34.5% of the fatty acids) and linolenic (~18% of fatty acids). These three fatty acids comprise over 90% of the fatty acids in CCWG-1-CuSO_4_. No long chain polyunsaturated long chain fatty acids such as EPA and DHA were detected. Results confirm that CCWG-1-CuSO_4_ contains negligible levels of erucic acid. Dairyland Laboratories did not detect it and EPL reported erucic acid to be less than 0.1% of the total fatty acids.

**Table 5.** CCWG-1-CuSO_4_ Fatty Acids (100% dry matter basis)

| Analytical |  |  | Standard |  |  | Range |  |
| --- | --- | --- | --- | --- | --- | --- | --- |
| Lab<sup>1</sup> | Variable | Units | Mean | Deviation | Median | Minimum | Maximum |
| EPL | Myristic (C14:0) | % | 0.02 | 0.001 | 0.02 | 0.02 | 0.02 |
| DLL | Myristic (C14:0) | % | 0.00 | 0.00 | 0.00 | 0.00 | 0.00 |
| EPL | Palmitic (C16:0) | % | 0.79 | 0.085 | 0.82 | 0.67 | 0.88 |
| DLL | Palmitic (C16:0) | % | 0.75 | 0.063 | 0.74 | 0.67 | 0.82 |
| EPL | Palmitoleic (C16:1) | % | 0.03 | 0.005 | 0.03 | 0.02 | 0.03 |
| DLL | Palmitoleic (C16:1) | % | 0.00 | 0.00 | 0.00 | 0.00 | 0.00 |
| EPL | Heptadecanoic (C17:0) | % | 0.01 | 0.001 | 0.01 | 0.01 | 0.01 |
| DLL | Heptadecanoic (C17:0) | % | 0.00 | 0.00 | 0.00 | 0.00 | 0.00 |
| EPL | Heptadecenoic (C17:1) | % | 0.01 | 0.001 | 0.01 | 0.01 | 0.02 |
| DLL | Heptadecenoic (C17:1) | % | NA<sup>2</sup> | NA | NA | NA | NA |
| EPL | Stearic (C18:0) | % | 0.20 | 0.024 | 0.19 | 0.17 | 0.23 |
| DLL | Stearic (C18:0) | % | 0.18 | 0.020 | 0.18 | 0.16 | 0.21 |
| EPL | Oleic (C18:1) | % | 8.04 | 0.962 | 8.04 | 6.73 | 9.19 |
| DLL | Oleic (C18:1) | % | 7.45 | 0.815 | 7.35 | 6.36 | 8.28 |
| EPL | Linoleic (C18:2) | % | 6.87 | 0.605 | 7.19 | 6.08 | 7.46 |
| DLL | Linoleic (C18:2) | % | 6.40 | 0.570 | 6.35 | 5.55 | 6.93 |
| EPL | Linolenic (C18:3) | % | 3.62 | 0.241 | 3.68 | 3.34 | 3.87 |
| DLL | Linolenic (C18:3) | % | 3.07 | 0.196 | 2.98 | 2.82 | 3.27 |
| EPL | Nonadecenoic (C19:1) | % | 0.01 | 0.001 | 0.01 | 0.01 | 0.01 |
| DLL | Nonadecenoic (C19:1) | % | NA | NA | NA | NA | NA |
| EPL | Arachidic (C20:0) | % | 0.03 | 0.004 | 0.03 | 0.03 | 0.04 |
| DLL | Arachidic (C20:0) | % | 0.00 | 0.00 | 0.00 | 0.00 | 0.00 |
| EPL | Eicosenoic (C20:1) | % | 0.16 | 0.011 | 0.16 | 0.15 | 0.17 |
| DLL | Eicosenoic (C20:1) | % | 0.00 | 0.00 | 0.00 | 0.00 | 0.00 |
| EPL | Eicosadienoic (C20:2) | % | 0.03 | 0.002 | 0.03 | 0.03 | 0.03 |
| DLL | Eicosadienoic (C20:2),<br>Eicosatrienoic(C20:3) | % | 0.45 | 0.179 | 0.51 | 0.17 | 0.64 |
| EPL | Behenic (C22:0) | % | 0.02 | 0.002 | 0.02 | 0.02 | 0.02 |
| DLL | Behenic (C22:0) | % | NA | NA | NA | NA | NA |
| EPL | Erucic (C22:1) | % | 0.06 | 0.013 | 0.07 | 0.04 | 0.08 |
| DLL | Erucic (C22:1) | % | 0.00 | 0.00 | 0.00 | 0.00 | 0.00 |
| EPL | Lignoceric (C24:0) | % | 0.02 | 0.002 | 0.02 | 0.01 | 0.02 |
| DLL | Lignoceric (C24:0) | % | 0.00 | 0.00 | 0.00 | 0.00 | 0.00 |
| EPL | Nervonic (C24:1) | % | 0.08 | 0.007 | 0.07 | 0.07 | 0.08 |
| DLL | Nervonic (C24:1) | % | 0.00 | 0.00 | 0.00 | 0.00 | 0.00 |

<sup>1</sup> Analytical Lab: EPL – EPL Bio Analytical Services; DLL – Dairyland Laboratories
<sup>2</sup> NA: Not Analyzed
The following fatty acids were below the level of detection for Lab EPL: Arachidonic (C20:4), Capric (C10:0), Caproic (C6:0), Caprylic (C8:0), Docosadienoic (C22:2), Eicosatrienoic (C20:3), Gamma Linolenic (C18:3), Heneicosanoic (C21:0), Lauric (C12:0), Myristoleic (C14:1), Nonadecanoic (C19:0), Pentadecanoic (C15:0), Pentadecenoic (C15:1), Tricosanoic (C23:0); for Lab DLL: Arachidonic (C20:4), Docosahexanoic acid (C22:6); Eicosapentaenoic acid (C20:5).

**Table 6.** CCWG-1-CuSO_4_ Fatty Acids as a Percent of Total Fatty Acids

| Analytical |  |  | Standard |  |  | Range |  |
| --- | --- | --- | --- | --- | --- | --- | --- |
| Lab<sup>1</sup> | Variable | Units | Mean | Deviation | Median | Minimum | Maximum |
| EPL | Myristic (C14:0) | % | 0.09 | 0.006 | 0.09 | 0.09 | 0.10 |
| DLL | Myristic (C14:0) | % | 0.00 | 0.00 | 0.00 | 0.00 | 0.00 |
| EPL | Palmitic (C16:0) | % | 3.98 | 0.066 | 4.00 | 3.86 | 4.02 |
| DLL | Palmitic (C16:0) | % | 4.06 | 0.076 | 4.04 | 4.00 | 4.20 |
| EPL | Palmitoleic (C16:1) | % | 0.15 | 0.012 | 0.16 | 0.13 | 0.16 |
| DLL | Palmitoleic (C16:1) | % | 0.00 | 0.00 | 0.00 | 0.00 | 0.00 |
| EPL | Heptadecanoic (C17:0) | % | 0.03 | 0.018 | 0.03 | 0.00 | 0.05 |
| DLL | Heptadecanoic (C17:0) | % | 0.00 | 0.00 | 0.00 | 0.00 | 0.00 |
| EPL | Heptadecenoic (C17:1) | % | 0.07 | 0.011 | 0.08 | 0.06 | 0.09 |
| DLL | Heptadecenoic (C17:1) | % | NA<sup>2</sup> | NA | NA | NA | NA |
| EPL | Stearic (C18:0) | % | 1.00 | 0.044 | 1.02 | 0.94 | 1.04 |
| DLL | Stearic (C18:0) | % | 0.99 | 0.050 | 1.01 | 0.91 | 1.03 |
| EPL | Oleic (C18:1) | % | 40.14 | 1.223 | 40.55 | 38.50 | 41.55 |
| DLL | Oleic (C18:1) | % | 40.07 | 0.943 | 40.19 | 38.61 | 41.06 |
| EPL | Linoleic (C18:2) | % | 34.41 | 0.611 | 34.20 | 33.80 | 35.30 |
| DLL | Linoleic (C18:2) | % | 34.48 | 0.337 | 34.39 | 34.17 | 34.94 |
| EPL | Linolenic (C18:3) | % | 18.15 | 0.722 | 17.90 | 17.55 | 19.40 |
| DLL | Linolenic (C18:3) | % | 16.55 | 0.632 | 16.32 | 16.18 | 17.67 |
| EPL | Nonadecenoic (C19:1) | % | 0.03 | 0.018 | 0.03 | 0.00 | 0.05 |
| DLL | Nonadecenoic (C19:1) | % | NA | NA | NA | NA | NA |
| EPL | Arachidic (C20:0) | % | 0.03 | 0.004 | 0.03 | 0.03 | 0.04 |
| DLL | Arachidic (C20:0) | % | 0.00 | 0.00 | 0.00 | 0.00 | 0.00 |
| EPL | Eicosenoic (C20:1) | % | 0.16 | 0.011 | 0.16 | 0.15 | 0.17 |
| DLL | Eicosenoic (C20:1) | % | 1.47 | 0.088 | 1.44 | 1.38 | 1.60 |
| EPL | Eicosadienoic (C20:2) | % | 0.03 | 0.002 | 0.03 | 0.03 | 0.03 |
| DLL | Eicosadienoic (C20:2),<br>Eicosatrienoic(C20:3) | % | 2.38 | 0.939 | 2.54 | 1.09 | 3.54 |
| EPL | Behenic (C22:0) | % | 0.02 | 0.002 | 0.02 | 0.02 | 0.02 |
| DLL | Behenic (C22:0) | % | 0.00 | 0.00 | 0.00 | 0.00 | 0.00 |
| EPL | Erucic (C22:1) | % | 0.06 | 0.013 | 0.07 | 0.04 | 0.08 |
| DLL | Erucic (C22:1) | % | 0.00 | 0.00 | 0.00 | 0.00 | 0.00 |
| EPL | Lignoceric (C24:0) | % | 0.07 | 0.021 | 0.08 | 0.04 | 0.09 |
| DLL | Lignoceric (C24:0) | % | 0.00 | 0.00 | 0.00 | 0.00 | 0.00 |
| EPL | Nervonic (C24:1) | % | 0.37 | 0.024 | 0.37 | 0.34 | 0.40 |
| DLL | Nervonic (C24:1) | % | 0.00 | 0.00 | 0.00 | 0.00 | 0.00 |

<sup>1</sup> Analytical Lab: EPL – EPL Bio Analytical Services; DLL – Dairyland Laboratories
<sup>2</sup> NA: Not Analyzed
The following fatty acids were below the level of detection for Lab A: Arachidonic (C20:4), Capric (C10:0), Caproic (C6:0), Caprylic (C8:0), Docosadienoic (C22:2), Eicosatrienoic (C20:3), Gamma Linolenic (C18:3), Heneicosanoic (C21:0), Lauric (C12:0), Myristoleic (C14:1), Nonadecanoic (C19:0), Pentadecanoic (C15:0), Pentadecenoic (C15:1), Tricosanoic (C23:0); for Lab B: Arachidonic (C20:4), Docosahexanoic acid (C22:6); Eicosapentaenoic acid (C20:5).

#### Glucosinolate Characterization and Quantification

The sole glucosinolate, sinigrin, averaged 85.7 and 86.9 μmoles/g of CCWG-1-CuSO_4_ for the two detection methods (Table 7). The median value was higher than the mean due to one lot containing almost half the amount of sinigrin as the other four lots. This is reflected by the low minimum value as compared to the maximum value. The much lower sinigrin level in the one lot may have resulted from the soil conditions at that site.

**Table 7.** CoverCress CS (CCWG-1-CuSO_4_) Glucosinolates (Sinigrin, 100% dry matter basis) – EPL Bio Analytical Services

| Variable | Units | Standard |  |  | Range |  |
| --- | --- | --- | --- | --- | --- | --- |
|  |  | Mean | Deviation | Median | Minimum | Maximum |
| Sinigrin – MS | $\mu\text{moles/g}$ | 85.7 | 16.83 | 90.7 | 56.2 | 98.6 |
| Sinigrin – UV | $\mu\text{moles/g}$ | 86.9 | 23.97 | 95.7 | 45 | 102 |

#### Minerals and Vitamins

Table 8 contains the results of the mineral analyses. EPL reported consistently higher mineral levels than Dairyland Laboratories, with the exception of zinc. The discrepancy between labs is unexplained. Generally, the values for the five lots had symmetric distribution. One location had a sulfur level that was 50% of the others. This lower sulfur value corresponds to the lower glucosinolate level of the sample from the same location. Glucosinolates are secondary sulfur compounds commonly found in brassica species (Aghajanzada *et al*., 2014), which suggests there is a plausible relationship between both values being lower at this site. The reason for the lower level at one location as compared to the other four locations is not clear but may have been a result of the soil fertilization. As expected, mean copper values were approximately 800 ppm because of the addition of copper sulfate. The analytical results confirm that the target level of 800 ppm copper was achieved.

**Table 8.**
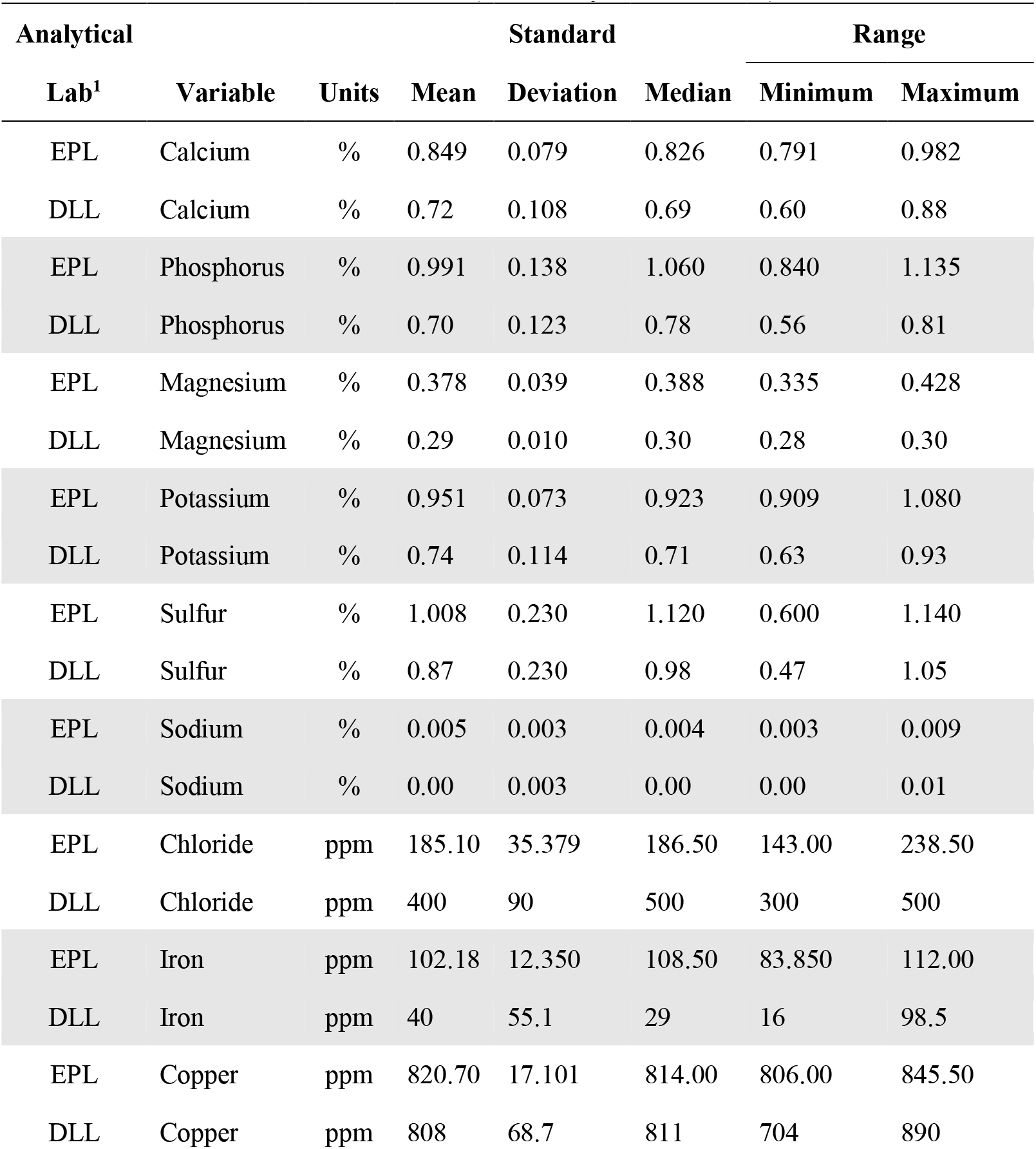
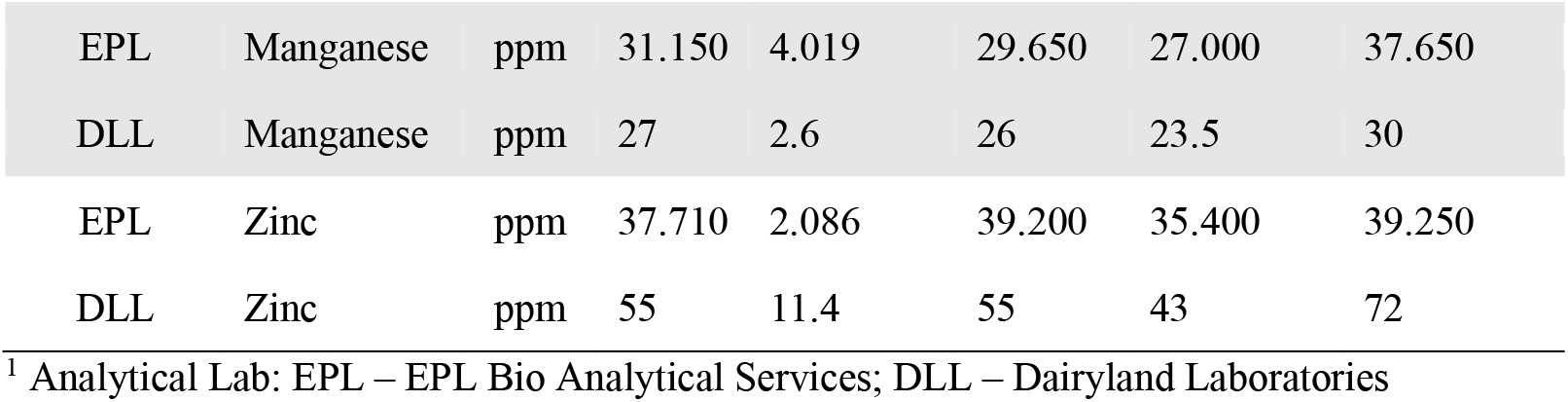
CCWG-1-CuSO_4_ Minerals (100% dry matter basis)

Table 9 contains the vitamins as analyzed by EPL for the five lots. Generally, the values for the five lots had symmetric distribution. Nutritionists and feed formulators, as common practice, do not use the vitamin content of feedstuffs in formulating diets due to the variability, stability, low levels in the feed ingredient and the low cost of supplementing vitamins. The Canola Council of Canada in their Canola Feed Guide (2019) state, “As is recommended with most natural sources of vitamins in animal feeds, users should not place too much reliance on these values and use supplemental vitamin premixes instead.”

**Table 9.** CCWG-1-CuSO_4_ Vitamins (100% dry matter basis) – EPL Bio Analytical Services

| Variable | Units | Standard |  |  | Range |  |
| --- | --- | --- | --- | --- | --- | --- |
|  |  | Mean | Deviation | Median | Minimum | Maximum |
| Folic Acid | mg/kg | 3.3 | 1.35 | 3.2 | 1.8 | 5.3 |
| Niacin | mg/kg | 51.2 | 10.52 | 48.6 | 42.1 | 69.3 |
| Vitamin B1 (Thiamine) | mg/kg | 5.4 | 0.25 | 5.3 | 5.2 | 5.8 |
| Vitamin B2 (Riboflavin) | mg/kg | 2.7 | 0.54 | 2.6 | 2.3 | 3.6 |
| Vitamin B6 (Pyridoxine) | mg/kg | 10.6 | 1.69 | 10.3 | 8.8 | 13.4 |
| Alpha-Tocopherol | mg/kg | 118.0 | 10.86 | 121.5 | 99.0 | 125.5 |
| Beta-Tocopherol | mg/kg | 1.8 | 0.07 | 1.8 | 1.7 | 1.9 |
| Delta-Tocopherol | mg/kg | 2.3 | 0.16 | 2.3 | 2.1 | 2.5 |
| Gamma-Tocopherol | mg/kg | 53.4 | 14.65 | 48.6 | 40.3 | 78.5 |

#### Sinapine and Mycotoxins

Sinapine, like glucosinolates, is an anti-nutrient found in modern oilseed rape and canola meals. Sinapine is metabolized in animals to trimethylamine, which is further metabolized by trimethyl oxidase. Certain brown egg laying strains of hens have been reported to have lower hepatic trimethyl oxidase activity apparently leading to accumulation of trimethylamine in some brown eggs and an associated “fishy taint”. As summarized by Rymer & Short (2003), sinapines are present in modern rapeseed at levels of 12-23 g/kg seed (Huisman & Tolman, 2001). They are converted in the large intestine to trimethylamine, which apparently produces an undesirable fishy odor in the eggs of certain brown egg chickens. Some of these types of birds lack the ability to produce trimethylamine oxidase (Jeroch, 2008), and as a result the trimethylamine accumulates in the eggs of these birds. This odor in eggs occurs at levels of 0.8 mg/kg diet (Fenwick, 1982). The concentrations of sinapine in CCWG-1-CuSO_4_ were <0.05% DM basis as reported by EPL in all five lots. This is much lower in comparison to canola meal where the Canola Council of Canada (2015) reports sinapine levels of 1.0% as is basis (with an average 12% moisture content) or 1.14% on a 100% DM basis.

The following mycotoxins were below the limits of detection in all samples from all of the five lots: Aflatoxin B1, B2, G1, G2; T-2 Toxin; Ochratoxin A; Deoxynivalenol (DON), 3-Acetyl-DON; 15-Acetyl-DON; Zearalenone; Fumonisin B1, B2, B3; Ergosine; Ergotamine; Ergocornine; and Ergocryptine. For Ergocristine, which is a natural ergot alkaloid, four lots were below the levels of detection. For one lot, one of the two replicates was below the level of detection and the second had 15.9 ppb on a dry matter basis with level of detection at 1.5 ppb.

### Study B – CCWG-2-CuSO_4_

#### Proximate Analyses

Table 10 contains the results of proximate analyses. Just as with the CCWG-1 samples, it was postulated that Dairyland Laboratories’s method extracted more fat from the product than EPL’s method. In addition, there was greater variation noted with EPL’s fat values than with Dairyland Labs. Fat in the sample may affect the fiber analysis with higher fat resulting in inflated crude fiber, ADF and NDF values. Moisture levels are high due to the addition of the copper sulfate solution to the CoverCress grain. Since the mean and median values are similar for all proximates, the values from the five lots are evenly distributed or symmetrical in distribution.

**Table 10.** CoverCress CS (CCWG-2-CuSO_4_) Proximates (100% dry matter basis)

| Analytical Lab<sup>1</sup> | Variable | Units | Standard |  |  | Range |  |
| --- | --- | --- | --- | --- | --- | --- | --- |
|  |  |  | Mean | Deviation | Median | Minimum | Maximum |
| EPL | Moisture | % | 19.6 | 0.24 | 19.6 | 19.3 | 19.9 |
| DLL | Moisture | % | 16.6 | 0.38 | 16.7 | 16.1 | 17.1 |
| EPL | Dry Matter | % | 80.4 | 0.24 | 80.5 | 80.1 | 80.7 |
| DLL | Dry Matter | % | 83.4 | 0.38 | 83.3 | 82.9 | 83.9 |
| EPL | Crude Fiber | % | 20.0 | 2.82 | 21.0 | 15.6 | 22.6 |
| DLL | Crude Fiber | % | 12.1 | 0.71 | 12.5 | 10.9 | 12.5 |
| EPL | Acid Detergent Fiber | % | 25.8 | 2.73 | 25.3 | 22.1 | 28.7 |
| DLL | Acid Detergent Fiber | % | 17.8 | 2.04 | 16.8 | 15.9 | 20.2 |
| EPL | Neutral Detergent Fiber | % | 26.9 | 3.32 | 27.7 | 22.0 | 30.2 |
| DLL | Neutral Detergent Fiber | % | 22.6 | 1.48 | 21.9 | 21.3 | 24.4 |
| EPL | Total Dietary Fiber | % | 28.8 | 1.04 | 28.7 | 27.7 | 30.3 |
| EPL | Crude Protein | % | 25.4 | 1.27 | 25.0 | 24.1 | 27.4 |
| DLL | Crude Protein | % | 23.9 | 0.97 | 24.1 | 22.5 | 25.1 |
| EPL | Crude Fat | % | 32.7 | 1.62 | 33.0 | 30.4 | 34.8 |
| DLL | Crude Fat | % | 37.0 | 1.07 | 37.0 | 35.7 | 38.5 |
| EPL | Carbohydrates | % | 36.3 | 2.12 | 36.8 | 32.7 | 38.1 |
| EPL | Ash | % | 5.6 | 0.45 | 5.4 | 5.1 | 6.2 |
| DLL | Ash | % | 6.0 | 0.86 | 6.1 | 4.6 | 6.7 |

<sup>1</sup> Analytical Lab: EPL – EPL Bio Analytical Services; DLL – Dairyland Laboratories

#### Amino Acid Analyses

Table 11 contains the amino acid profile of CCWG-2-CuSO_4_ when expressed on a 100% dry matter basis and Table 12 contains the amino acid profile of CCWG-2-CuSO_4_ when expressed on a protein basis. The amino acid values from the five lots showed a symmetrical distribution with the means and median values being similar. Dairyland had a higher percentage of total amino acids when total amino acids were expressed as a percent of crude protein (88.1% versus 83.1%).

**Table 11.** CoverCress CS (CCWG-2-CuSO_4_) Amino Acids (100% dry matter basis)

| Analytical Lab | Variable | Units | Mean | Standard Deviation | Median | Range |  |
| --- | --- | --- | --- | --- | --- | --- | --- |
|  |  |  |  |  |  | Minimum | Maximum |
| EPL | Alanine | % | 1.13 | 0.023 | 1.14 | 1.11 | 1.15 |
| DLL | Alanine | % | 1.00 | 0.034 | 0.99 | 0.97 | 1.06 |
| EPL | Arginine | % | 1.39 | 0.067 | 1.39 | 1.30 | 1.46 |
| DLL | Arginine | % | 1.57 | 0.052 | 1.55 | 1.51 | 1.63 |
| EPL | Aspartic Acid | % | 1.93 | 0.054 | 1.91 | 1.91 | 2.03 |
| DLL | Aspartic Acid | % | 1.92 | 0.031 | 1.93 | 1.88 | 1.96 |
| EPL | Cystine | % | 0.38 | 0.022 | 0.38 | 0.36 | 0.41 |
| DLL | Cysteine | % | 0.38 | 0.017 | 0.39 | 0.36 | 0.41 |
| EPL | Glutamic Acid | % | 3.83 | 0.110 | 3.81 | 3.68 | 3.95 |
| DLL | Glutamic Acid | % | 3.59 | 0.133 | 3.59 | 3.40 | 3.75 |
| EPL | Glycine | % | 1.38 | 0.064 | 1.38 | 1.30 | 1.46 |
| DLL | Glycine | % | 1.45 | 0.041 | 1.45 | 1.39 | 1.51 |
| EPL | Histidine | % | 0.54 | 0.022 | 0.54 | 0.52 | 0.57 |
| DLL | Histidine | % | 0.47 | 0.015 | 0.47 | 0.45 | 0.49 |
| EPL | Isoleucine | % | 0.93 | 0.019 | 0.93 | 0.90 | 0.94 |
| DLL | Isoleucine | % | 0.73 | 0.047 | 0.74 | 0.68 | 0.80 |
| EPL | Leucine | % | 1.63 | 0.046 | 1.63 | 1.56 | 1.68 |
| DLL | Leucine | % | 1.59 | 0.059 | 1.58 | 1.53 | 1.68 |
| EPL | Lysine | % | 1.28 | 0.035 | 1.28 | 1.24 | 1.34 |
| DLL | Lysine | % | 1.12 | 0.032 | 1.13 | 1.09 | 1.17 |
| EPL | Methionine | % | 0.37 | 0.029 | 0.38 | 0.33 | 0.41 |
| DLL | Methionine | % | 0.33 | 0.019 | 0.33 | 0.31 | 0.36 |
| EPL | Phenylalanine | % | 0.99 | 0.060 | 0.98 | 0.92 | 1.08 |
| DLL | Phenylalanine | % | 1.04 | 0.033 | 1.04 | 0.99 | 1.08 |
| EPL | Proline | % | 1.22 | 0.043 | 1.23 | 1.17 | 1.27 |
| DLL | Proline | % | 1.36 | 0.020 | 1.37 | 1.33 | 1.38 |
| EPL | Serine | % | 0.93 | 0.035 | 0.93 | 0.89 | 0.99 |
| DLL | Serine | % | 0.99 | 0.019 | 0.99 | 0.97 | 1.02 |
| EPL | Threonine | % | 1.03 | 0.030 | 1.04 | 0.99 | 1.07 |
| DLL | Threonine | % | 1.06 | 0.025 | 1.05 | 1.03 | 1.10 |
| EPL | Tryptophan | % | 0.36 | 0.016 | 0.37 | 0.33 | 0.37 |
| DLL | Tryptophan | % | 0.47 | 0.059 | 0.47 | 0.42 | 0.56 |
| EPL | Tyrosine | % | 0.49 | 0.032 | 0.49 | 0.45 | 0.54 |
| DLL | Tyrosine | % | 0.74 | 0.018 | 0.74 | 0.72 | 0.75 |
| EPL | Valine | % | 1.23 | 0.019 | 1.23 | 1.20 | 1.26 |
| DLL | Valine | % | 1.17 | 0.063 | 1.17 | 1.11 | 1.27 |

<sup>1</sup> Analytical Lab: EPL – EPL Bio Analytical Services; DLL – Dairyland Laboratories

**Table 12.** CoverCress CS (CCWG-2-CuSO_4_) Amino Acids as Percent of Protein

| Analytical Lab | Variable | Units | Mean | Standard Deviation | Median | Range |  |
| --- | --- | --- | --- | --- | --- | --- | --- |
|  |  |  |  |  |  | Minimum | Maximum |
| EPL | Alanine | % | 4.46 | 0.164 | 4.48 | 4.20 | 4.60 |
| DLL | Alanine | % | 4.20 | 0.118 | 4.19 | 4.05 | 4.38 |
| EPL | Arginine | % | 5.49 | 0.160 | 5.44 | 5.31 | 5.70 |
| DLL | Arginine | % | 6.59 | 0.138 | 6.59 | 6.43 | 6.74 |
| EPL | Aspartic Acid | % | 7.64 | 0.450 | 7.73 | 6.97 | 8.12 |
| DLL | Aspartic Acid | % | 8.05 | 0.220 | 7.99 | 7.81 | 8.33 |
| EPL | Cystine | % | 1.50 | 0.104 | 1.50 | 1.41 | 1.67 |
| DLL | Cysteine | % | 1.62 | 0.076 | 1.62 | 1.49 | 1.69 |
| EPL | Glutamic Acid | % | 15.13 | 0.711 | 15.35 | 13.90 | 15.75 |
| DLL | Glutamic Acid | % | 15.07 | 0.172 | 15.09 | 14.88 | 15.27 |
| EPL | Glycine | % | 5.45 | 0.094 | 5.41 | 5.36 | 5.56 |
| DLL | Glycine | % | 6.09 | 0.087 | 6.06 | 6.01 | 6.20 |
| EPL | Histidine | % | 2.14 | 0.043 | 2.14 | 2.08 | 2.19 |
| DLL | Histidine | % | 1.99 | 0.027 | 1.99 | 1.96 | 2.02 |
| EPL | Isoleucine | % | 3.66 | 0.124 | 3.72 | 3.45 | 3.74 |
| DLL | Isoleucine | % | 3.07 | 0.125 | 3.09 | 2.87 | 3.19 |
| EPL | Leucine | % | 6.42 | 0.162 | 6.50 | 6.13 | 6.51 |
| DLL | Leucine | % | 6.69 | 0.081 | 6.71 | 6.56 | 6.78 |
| EPL | Lysine | % | 5.05 | 0.330 | 5.08 | 4.53 | 5.35 |
| DLL | Lysine | % | 4.71 | 0.121 | 4.69 | 4.55 | 4.85 |
| EPL | Methionine | % | 1.47 | 0.083 | 1.49 | 1.38 | 1.57 |
| DLL | Methionine | % | 1.39 | 0.050 | 1.38 | 1.33 | 1.45 |
| EPL | Phenylalanine | % | 3.89 | 0.072 | 3.91 | 3.80 | 3.96 |
| DLL | Phenylalanine | % | 4.35 | 0.044 | 4.38 | 4.30 | 4.40 |
| EPL | Proline | % | 4.83 | 0.136 | 4.87 | 4.60 | 4.95 |
| DLL | Proline | % | 5.73 | 0.164 | 5.71 | 5.48 | 5.91 |
| EPL | Serine | % | 3.68 | 0.062 | 3.69 | 3.60 | 3.76 |
| DLL | Serine | % | 4.16 | 0.116 | 4.13 | 4.06 | 4.36 |
| EPL | Threonine | % | 4.08 | 0.103 | 4.11 | 3.91 | 4.18 |
| DLL | Threonine | % | 4.44 | 0.098 | 4.38 | 4.36 | 4.58 |
| EPL | Tryptophan | % | 1.41 | 0.058 | 1.44 | 1.34 | 1.48 |
| DLL | Tryptophan | % | 1.98 | 0.174 | 1.93 | 1.80 | 2.21 |
| EPL | Tyrosine | % | 1.93 | 0.051 | 1.97 | 1.87 | 1.97 |
| DLL | Tyrosine | % | 3.09 | 0.073 | 3.08 | 2.99 | 3.18 |
| EPL | Valine | % | 4.85 | 0.170 | 4.92 | 4.56 | 4.98 |
| DLL | Valine | % | 4.89 | 0.180 | 4.98 | 4.59 | 5.04 |

<sup>1</sup> Analytical Lab: EPL – EPL Bio Analytical Services; DLL – Dairyland Laboratories

#### Fatty Acid Characterization

Tables 13 and 14 contains the fatty acid profile of CCWG-2-CuSO_4_ when expressed on a 100% dry matter basis and percent of total fatty acids, respectively. In general, there was good agreement between laboratories. The fatty acid values from the five lots showed a symmetrical distribution with the means and median values being similar. The major fatty acids are oleic (~43% of the total fatty acids) followed by linoleic (~32% of the total fatty acids) and linolenic (~18% of the total fatty acids). These three fatty acids comprise over 90% of the fatty acids in CCWG-2-CuSO_4_. No high polyunsaturated long chain fatty acids such as EPA and DHA were detected. Results confirm that CCWG-2-CuSO_4_ contains negligible levels of erucic acid. Dairyland Laboratories did not detect it and EPL reported erucic acid to be less than 0.5% of the total fatty acids.

**Table 13.** CoverCress CS (CCWG-2-CuSO_4_) Fatty Acids (100% dry matter basis)

| Analytical Lab | Variable | Units | Mean | Standard Deviation | Median | Range |  |
| --- | --- | --- | --- | --- | --- | --- | --- |
|  |  |  |  |  |  | Minimum | Maximum |
| EPL | Myristic (C14:0) | % | 0.02 | 0.002 | 0.02 | 0.01 | 0.02 |
| DLL | Myristic (C14:0) | % | 0.00 | 0.000 | 0.00 | 0.00 | 0.00 |
| EPL | Palmitic (C16:0) | % | 0.62 | 0.039 | 0.62 | 0.56 | 0.67 |
| DLL | Palmitic (C16:0) | % | 1.17 | 0.051 | 1.16 | 1.11 | 1.24 |
| EPL | Palmitoleic (C16:1) | % | 0.02 | 0.002 | 0.02 | 0.02 | 0.03 |
| DLL | Palmitoleic (C16:1) | % | 0.00 | 0.000 | 0.00 | 0.00 | 0.00 |
| EPL | Heptadecanoic (C17:0) | % | 0.01 | 0.001 | 0.01 | 0.01 | 0.01 |
| DLL | Heptadecanoic (C17:0) | % | 0.00 | 0.000 | 0.00 | 0.00 | 0.00 |
| EPL | Heptadecenoic (C17:1) | % | 0.00 | 0.000 | 0.00 | 0.00 | 0.00 |
| DLL | Heptadecenoic (C17:1) | % | NA<sup>2</sup> | NA | NA | NA | NA |
| EPL | Stearic (C18:0) | % | 0.13 | 0.010 | 0.13 | 0.12 | 0.14 |
| DLL | Stearic (C18:0) | % | 0.26 | 0.017 | 0.27 | 0.24 | 0.28 |
| EPL | Oleic (C18:1) | % | 6.75 | 0.430 | 6.74 | 6.17 | 7.38 |
| DLL | Oleic (C18:1) | % | 13.61 | 0.930 | 13.84 | 12.35 | 14.86 |
| EPL | Linoleic (C18:2) | % | 5.16 | 0.411 | 5.14 | 4.77 | 5.84 |
| DLL | Linoleic (C18:2) | % | 9.57 | 0.733 | 9.83 | 8.57 | 10.44 |
| EPL | Linolenic (C18:3) | % | 2.91 | 0.250 | 2.83 | 2.68 | 3.34 |
| DLL | Linolenic (C18:3) | % | 5.16 | 0.423 | 5.28 | 4.72 | 5.70 |
| EPL | Nonadecenoic (C19:1) | % | 0.00 | 0.000 | 0.00 | 0.00 | 0.00 |
| DLL | Nonadecenoic (C19:1) | % | NA | NA | NA | NA | NA |
| EPL | Arachidic (C20:0) | % | 0.02 | 0.001 | 0.02 | 0.02 | 0.02 |
| DLL | Arachidic (C20:0) | % | 0.00 | 0.000 | 0.00 | 0.00 | 0.00 |
| EPL | Eicosenoic (C20:1) | % | 0.14 | 0.009 | 0.14 | 0.12 | 0.14 |
| DLL | Eicosenoic (C20:1) | % | 0.23 | 0.014 | 0.23 | 0.21 | 0.25 |
| EPL | Eicosadienoic (C20:2) | % | 0.02 | 0.003 | 0.02 | 0.02 | 0.03 |
| DLL | Eicosadienoic (C20:2),<br>Eicosatrienoic(C20:3) | % | 0.00 | 0.000 | 0.00 | 0.00 | 0.00 |
| EPL | Behenic (C22:0) | % | 0.00 | 0.000 | 0.00 | 0.00 | 0.00 |
| DLL | Behenic (C22:0) | % | NA | NA | NA | NA | NA |
| EPL | Erucic (C22:1) | % | 0.07 | 0.022 | 0.06 | 0.05 | 0.11 |
| DLL | Erucic (C22:1) | % | 0.00 | 0.000 | 0.00 | 0.00 | 0.00 |
| EPL | Lignoceric (C24:0) | % | 0.00 | 0.000 | 0.00 | 0.00 | 0.00 |
| DLL | Lignoceric (C24:0) | % | 0.00 | 0.000 | 0.00 | 0.00 | 0.00 |
| EPL | Nervonic (C24:1) | % | 0.05 | 0.005 | 0.05 | 0.04 | 0.06 |
| DLL | Nervonic (C24:1) | % | 0.00 | 0.00 | 0.00 | 0.00 | 0.00 |

<sup>1</sup> Analytical Lab: EPL – EPL Bio Analytical Services; DLL – Dairyland Laboratories
<sup>2</sup> NA: Not Analyzed
The following fatty acids were below the level of detection for Lab EPL: Capric (C10:0), Caproic (C6:0), Caprylic (C8:0), Docosadienoic (C22:2), Docosahexaenoic (C22:6), Eicosatrienoic (C20:3), Eicosapentaenoic (C20:5), Gamma Linolenic (C18:3), Heneicosanoic (C21:0), Heptadecenoic (C17:1), Lauric (C12:0), Myristoleic (C14:1), Nonadecanoic (C19:0), Nonadecenoic (C19:1), Pentadecanoic (C15:0), Pentadecenoic (C15:1), Tricosanoic (C23:0); for Lab DLL: Arachidonic (C20:4), Docosahexanoic acid (C22:6); Eicosapentaenoic acid (C20:5).

**Table 14.** CoverCress CS (CCWG-2-CuSO_4_) Fatty Acids as a Percent of Total Fatty Acids

| Analytical<br>Lab | Variable | Units | Mean | Standard<br>Deviation | Median | Range |  |
| --- | --- | --- | --- | --- | --- | --- | --- |
|  |  |  |  |  |  | Minimum | Maximum |
| EPL | Myristic (C14:0) | % | 0.10 | 0.011 | 0.10 | 0.09 | 0.11 |
| DLL | Myristic (C14:0) | % | 0.00 | 0.00 | 0.00 | 0.00 | 0.00 |
| EPL | Palmitic (C16:0) | % | 3.88 | 0.042 | 3.89 | 3.84 | 3.95 |
| DLL | Palmitic (C16:0) | % | 3.89 | 0.079 | 3.91 | 3.77 | 3.96 |
| EPL | Palmitoleic (C16:1) | % | 0.15 | 0.018 | 0.16 | 0.12 | 0.17 |
| DLL | Palmitoleic (C16:1) | % | 0.00 | 0.00 | 0.00 | 0.00 | 0.00 |
| EPL | Heptadecanoic (C17:0) | % | 0.03 | 0.018 | 0.03 | 0.00 | 0.05 |
| DLL | Heptadecanoic (C17:0) | % | 0.00 | 0.00 | 0.00 | 0.00 | 0.00 |
| EPL | Heptadecenoic (C17:1) | % | 0.07 | 0.011 | 0.08 | 0.06 | 0.09 |
| DLL | Heptadecenoic (C17:1) | % | NA | NA | NA | NA | NA |
| EPL | Stearic (C18:0) | % | 0.84 | 0.039 | 0.84 | 0.77 | 0.88 |
| DLL | Stearic (C18:0) | % | 0.88 | 0.040 | 0.87 | 0.83 | 0.93 |
| EPL | Oleic (C18:1) | % | 42.32 | 2.261 | 42.40 | 39.00 | 45.35 |
| DLL | Oleic (C18:1) | % | 45.40 | 2.134 | 45.34 | 42.22 | 48.16 |
| EPL | Linoleic (C18:2) | % | 32.42 | 1.425 | 32.50 | 30.25 | 34.25 |
| DLL | Linoleic (C18:2) | % | 31.89 | 1.431 | 31.86 | 29.84 | 33.87 |
| EPL | Linolenic (C18:3) | % | 18.25 | 0.793 | 18.15 | 17.35 | 19.50 |
| DLL | Linolenic (C18:3) | % | 17.18 | 0.737 | 17.24 | 16.44 | 18.26 |
| EPL | Nonadecenoic (C19:1) | % | 0.03 | 0.018 | 0.03 | 0.00 | 0.05 |
| DLL | Nonadecenoic (C19:1) | % | NA | NA | NA | NA | NA |
| EPL | Arachidic (C20:0) | % | 0.13 | 0.013 | 0.13 | 0.13 | 0.016 |
| DLL | Arachidic (C20:0) | % | 0.00 | 0.00 | 0.00 | 0.00 | 0.00 |
| EPL | Eicosenoic (C20:1) | % | 0.85 | 0.024 | 0.84 | 0.82 | 0.87 |
| DLL | Eicosenoic (C20:1) | % | 0.77 | 0.042 | 0.78 | 0.70 | 0.82 |
| EPL | Eicosadienoic (C20:2) | % | 0.03 | 0.002 | 0.03 | 0.03 | 0.03 |
| DLL | Eicosadienoic (C20:2),<br>Eicosatrienoic(C20:3) | % | 0.00 | 0.000 | 0.00 | 0.00 | 0.00 |
| DLL | Arachidonic (20:4) | % | 0.02 | 0.018 | 0.02 | 0.00 | 0.05 |
| EPL | Behenic (C22:0) | % | 0.06 | 0.003 | 0.06 | 0.06 | 0.07 |
| DLL | Behenic (C22:0) | % | 0.00 | 0.00 | 0.00 | 0.00 | 0.00 |
| EPL | Erucic (C22:1) | % | 0.43 | 0.109 | 0.38 | 0.36 | 0.62 |
| DLL | Erucic (C22:1) | % | 0.00 | 0.00 | 0.00 | 0.00 | 0.00 |
| EPL | Lignoceric (C24:0) | % | 0.07 | 0.005 | 0.07 | 0.06 | 0.07 |
| DLL | Lignoceric (C24:0) | % | 0.00 | 0.00 | 0.00 | 0.00 | 0.00 |
| EPL | Nervonic (C24:1) | % | 0.31 | 0.025 | 0.30 | 0.30 | 0.36 |
| DLL | Nervonic (C24:1) | % | 0.00 | 0.00 | 0.00 | 0.00 | 0.00 |

<sup>1</sup> Analytical Lab: EPL – EPL Bio Analytical Services; DLL – Dairyland Laboratories
<sup>2</sup> NA: Not Analyzed
The following fatty acids were below the level of detection for Lab A: Arachidonic (C20:4), Capric (C10:0), Caproic (C6:0), Caprylic (C8:0), Docosadienoic (C22:2), Eicosatrienoic (C20:3), Gamma Linolenic (C18:3), Heneicosanoic (C21:0), Lauric (C12:0), Myristoleic (C14:1), Nonadecanoic (C19:0), Pentadecanoic (C15:0), Pentadecenoic (C15:1), Tricosanoic (C23:0); for Lab B: Arachidonic (C20:4), Docosahexanoic acid (C22:6); Eicosapentaenoic acid (C20:5).

#### Glucosinolate Characterization and Quantification

Sinigrin averaged 105.4 μmoles/g of CCWG-2-CuSO_4_ on a 100% dry matter basis and was similar among lots (Table 15). Sinigrin was measured using the Ultraviolet method (UV).

**Table 15.** CoverCress CS (CCWG-2-CuSO_4_) Glucosinolates (Sinigrin, 100% dry matter basis) - EPL Bio Analytical Services

| Variable | Units | Mean | Standard<br>Deviation | Median | Range |  |
| --- | --- | --- | --- | --- | --- | --- |
|  |  |  |  |  | Minimum | Maximum |
| Sinigrin - UV | μmoles/g | 105.4 | 4.21 | 104.3 | 101.4 | 110.7 |

#### Minerals, Sinapine, and Mycotoxins

Table 16 contains the results to the mineral analyses. EPL and Dairyland Laboratories results varied. The discrepancy between labs is unexplained. Generally, the values for the five lots had symmetric distribution. The high level of copper is due to the addition of copper sulfate. Target addition of copper was 800 ppm. The analytical results confirm that the target levels were achieved.

**Table 16.** CoverCress CS (CCWG-2-CuSO_4_) Minerals (100% dry matter basis)

| Analytical Lab<sup>1</sup> | Variable | Units | Mean | Standard |  | Range |  |
| --- | --- | --- | --- | --- | --- | --- | --- |
|  |  |  |  | Deviation | Median | Minimum | Maximum |
| EPL | Calcium | % | 0.93 | 0.074 | 0.95 | 0.84 | 1.01 |
| DLL | Calcium | % | 1.25 | 0.154 | 1.31 | 0.99 | 1.39 |
| EPL | Phosphorus | % | 0.99 | 0.135 | 0.96 | 0.80 | 1.14 |
| DLL | Phosphorus | % | 0.68 | 0.134 | 0.71 | 0.48 | 0.81 |
| EPL | Magnesium | % | 0.38 | 0.091 | 0.37 | 0.30 | 0.48 |
| DLL | Magnesium | % | 0.35 | 0.087 | 0.30 | 0.27 | 0.44 |
| EPL | Potassium | % | 1.11 | 0.083 | 1.05 | 1.05 | 1.23 |
| DLL | Potassium | % | 1.04 | 0.117 | 1.05 | 0.89 | 1.17 |
| EPL | Sulfur | % | 1.17 | 0.061 | 1.15 | 1.12 | 1.26 |
| DLL | Sulfur | % | 0.94 | 0.064 | 0.96 | 0.86 | 1.01 |
| EPL | Sodium | % | 0.00 | 0.000 | 0.00 | 0.00 | 0.00 |
| DLL | Sodium | % | 0.01 | 0.004 | 0.01 | 0.01 | 0.02 |
| EPL | Chloride | ppm | 399.4 | 85.334 | 432.00 | 275.50 | 473.00 |
| DLL | Chloride | ppm | 800 | 25 | 700 | 600 | 1200 |
| EPL | Iron | ppm | 107.85 | 48.471 | 84.95 | 63.90 | 162.50 |
| DLL | Iron | ppm | 72 | 30.7 | 57 | 40.5 | 113 |
| EPL | Copper | ppm | 959.70 | 29.641 | 955.00 | 930.50 | 1001.5 |
| DLL | Copper | ppm | 1052 | 257.8 | 1087.5 | 633 | 1315 |
| EPL | Manganese | ppm | 36.96 | 3.254 | 38.30 | 31.80 | 39.75 |
| DLL | Manganese | ppm | 46 | 6.5 | 47.5 | 39 | 55 |
| EPL | Zinc | ppm | 38.88 | 4.672 | 40.20 | 33.65 | 45.00 |
| DLL | Zinc | ppm | 46 | 15.6 | 43 | 34 | 73 |

<sup>1</sup> Analytical Lab: EPL – EPL Bio Analytical Services; DLL – Dairyland Laboratories

The concentrations of sinapine in CCWG-2-CuSO_4_ were <0.05% as reported by EPL in all five lots. This is in comparison to canola meal where the Canola Council of Canada (2015) reports sinapine levels of 1.0% as is basis (with an average 12% moisture content) or 1.14% on a 100% DM basis.

No mycotoxins were detected in the five lots. All mycotoxins were below the limits of detection in all samples from all of the five lots: Aflatoxin B1, B2, G1, G2; T-2 Toxin; Ochratoxin A; Deoxynivalenol (DON), 3-Acetyl-DON; 15-Acetyl-DON; Zearalenone; Fumonisin B1, B2, B3; Ergosine; Ergotamine; Ergocristine; and Ergocornine; and Ergocryptine.

#### Vitamins

Table 17 contains the vitamins as analyzed by EPL for the five lots. Generally, the values for the five lots had symmetric distribution.

**Table 17.** CoverCress CS (CCWG-2-CuSO_4_) Vitamins (100% dry matter basis) – EPL Bio Analytical Services

| Variable | Units | Mean | Standard Deviation | Median | Range |  |
| --- | --- | --- | --- | --- | --- | --- |
|  |  |  |  |  | Minimum | Maximum |
| Folic Acid | mg/kg | 5.6 | 0.67 | 5.2 | 5.0 | 6.6 |
| Niacin | mg/kg | 40.2 | 2.06 | 40.6 | 37.1 | 42.5 |
| Vitamin B1 (Thiamine) | mg/kg | 5.2 | 0.06 | 5.2 | 5.1 | 5.3 |
| Vitamin B2 (Riboflavin) | mg/kg | 4.7 | 0.29 | 4.6 | 4.3 | 5.0 |
| Vitamin B6 (Pyridoxine) | mg/kg | 6.1 | 0.17 | 6.1 | 5.9 | 6.3 |
| Alpha-Tocopherol | mg/kg | 181.8 | 24.28 | 174.5 | 160.5 | 218.0 |
| Beta-Tocopherol | mg/kg | 3.1 | 0.13 | 3.1 | 2.9 | 3.4 |
| Delta-Tocopherol | mg/kg | <1.25 |  | <1.25 | <1.25 | <1.25 |
| Gamma-Tocopherol | mg/kg | 13.4 | 1.75 | 12.5 | 11.9 | 15.8 |

## Conclusions

The results of these extensive compositional and nutritional analyses show:

- The composition and nutritional components of CCWG-1-CuSO_4_ and CCWG-2-CuSO_4_ lots were generally consistent with very good process control and low inter-lot variability
- As expected by loss of function of the FAE1 gene, the fatty acid profile of the CCWG-1-CuSO_4_ and CCWG-2-CuSO_4_ lots showed negligible erucic acid levels
- As expected by loss of function of the TT8 gene, mean ADF and crude fibers levels were in acceptable ranges for broiler diets
- Sinigrin (2-propenyl glucosinolate) was the only detectable glucosinolate consistent with previous published results in field pennycress seed or meals and total glucosinolate concentration in CCWG-1-CuSO_4_ ranged from 45 to 102 μmoles/g on 100% DM basis. Total glucosinolate concentration in CCWG-2-CuSO_4_ ranged from 101.4 to 110.7 μmoles/g on 100% DM basis
- Total copper levels achieved target levels of 800 ppm
- Other anti-nutrients (sinapine) and mycotoxins were below the imit of detection or quantification

The main purpose of CoverCress whole grain as a feed ingredient in animal diets (e.g. broilers) is as an energy source. Determination of energy value is complex, but is related to the fat, protein, and fiber content. Further, the degree of unsaturation of the fatty acids is a factor in energy. While there are some slight differences in total fat and the fiber profile, the overall nutritional value of CCWG for energy is expected to be similar to that of canola seed (see Table 18 below). Levels of the four key limiting amino acids in broiler diets, methionine, lysine, threonine and valine are similar between CCWG and canola. The low-erucic acid phenotype expected with loss of function of FAE1 was achieved as shown by the low to non-detectable levels in CCWG. The nutritional value in terms of energy (total fat, profile of fatty acids, crude fiber, ADF, NDF) of CCWG-2 is comparable to that of CCWG-1. Crude fiber levels were variable across lots and between CCWG-1 and CCWG-2. This may be due to assay or natural variability within the plants. Levels of the four key limiting amino acids in broiler diets, methionine, lysine, threonine and valine were lower for the CCWG-2 as compared to the CCWG-1. The difference is mostly like due to the lower amino acids as a percent of protein in the CCWG-2 as compared to the CCWG-1 (88.1% versus 97% of the protein for CCWG-2 and CCWG-1, respectively.

**Table 18.** Key Compositional Parameters for Canola Whole Grain, CCWG-1 and CCWG-2

| Component | Canola<sup>1</sup> | CCWG-1-CuSO<sub>4</sub><br>Mean<sup>2</sup><br>[range] | CCWG-2-CuSO<sub>4</sub><br>Mean<sup>2</sup><br>[range] |
| --- | --- | --- | --- |
| Fat % DM | 39.1 | 36.3<br>[35.5-36.7] | 37.0<br>[35.7-38.5] |
| Erucic acid<br>% TFA | <0.1 | <0.1 | <0.1 |
| Oleic % of TFA | 61.4 | 40.07<br>[38.61-41.06] | 45.4<br>[42.22-48.16] |
| Linoleic % TFA | 20.1 | 34.48<br>[34.17-34.94] | 31.89<br>[29.84-33.87] |
| Linolenic % TFA | 9.3 | 16.55<br>[16.18-17.67] | 17.18<br>[16.44-18.26] |
| Total saturated %TFA | 7.0 | 5.0 |  |
| Total mono-unsaturated %TFA | 64.4 | 41.6 |  |
| Total poly-unsaturated % TFA | 28.6 | 53.4 |  |
| Protein %DM | 24.5 | 25.5<br>[22.7-26.7] | 23.9<br>[22.5-25.1] |
| Methionine % protein | 1.93 | 1.88<br>[1.74-2.03] | 1.39<br>[1.33-1.45] |
| Lysine % protein | 5.66 | 5.51<br>[5.30-5.68] | 4.71<br>[4.55-4.85] |
| Threonine % protein | 3.97 | 4.91<br>[4.62-5.26] | 4.44<br>[4.36-4.58] |
| Valine %protein | 4.40 | 5.69<br>[5.48-6.14] | 4.89<br>[4.59-5.04] |
| Crude Fiber %DM | 13.3 | 23.0<br>[20.9-24.9] | 12.1<br>[10.9-12.5] |
| ADF | 18.1 | 13.9<br>[12.1-15.8] | 17.8<br>[15.9-20.2] |
| NDF | 25.7 | 20.3<br>[17.3-24.5] | 22.6<br>[21.3-24.4] |
| Sulfur % DM | 0.45<sup>3</sup><br>[0.36-0.55] | 0.87<br>[0.47-1.05] | 0.94<br>[0.86-1.01] |
| Total glucosinolates µmoles/g on DM basis UV method | 9.8 | 86.9<br>[45-102] | 105.4<br>[101.4-110.7] |

<sup>1</sup>Values from Dairy One feed Composition Laboratory; means of whole canola seed samples from crop years 2004 - 2020. https://www.dairyoneservices.com/feedcomposition/
<sup>2</sup>From the respective 5-lot study, DLL values
<sup>3</sup>From https://www.grainscanada.gc.ca/en/grain-research/export-quality/oilseeds/canola/2020/index.htm

The composition and nutritional components of CCWG-1-CuSO4 and CCWG-2-CuSO_4_ lots were generally consistent with very good process control and inter-lot variability. These results will enable eventual development of permissible ranges for key nutritional analytes and guaranteed levels for specific components (i.e. commercial specifications) like total crude protein, crude fiber, crude fat, copper and sulfur. A glucosinolate specification will be needed to ensure safe consumption depending on the level of inclusion in the diet and sensitivity of the animal species. In these lots, the mean level of glucosinolates was slightly higher in CCWG-2 than CCWG-1, but ranges for the two overlapped, indicating the difference is likely due to natural variability.

The current study shows that low-erucic acid, lower-fiber pennycress (CoverCress) produces a consistent compositional phenotype that may enable the seed to be consumed as an energy source for various animal species.

## Acknowledgements

The authors would like to acknowledge Shengjun Liu (CoverCress Inc; St. Louis, MO) for treatment of the CoverCress grain with copper sulfate solution, Margaret Nemeth (Statistical Consultants Plus; Fenton, MO) for statistical compilation and CoverCress Inc. (St. Louis MO) for financial support.

## Footnotes

1 For simplicity, CoverCress is not marked with ™ for the remainder of the document.

